# Epigenetic analysis identifies factors driving racial disparity in prostate cancer

**DOI:** 10.1101/352138

**Authors:** Richa Rai, Shalini S Yadav, Heng Pan, James O’Connor, Mohamed Alshalalfa, Elai Davicioni, Emanuela Taioli, Olivier Elemento, Ashutosh K Tewari, Kamlesh K Yadav

## Abstract

Prostate cancer (PCa) is the second most leading cause of death in men worldwide. African American men (AA) represent more aggressive form of PCa as compared to Caucasian (CA) counterparts. Evidence suggests that genetic and other biological factors could account for the observed racial disparity. We analyzed the cancer genome atlas (TCGA) dataset (2015) for existing epigenetic variation in AA and CA prostate cancer patients, and carried out Reduced Representation Bisulphite Sequencing (RRBS) analysis to identify global methylation changes in AA and CA prostate cancer patients. The TCGA dataset analysis revealed that the epigenetic heterogeneity could be categorized into 4 classes, where AA associated primarily to methylation cluster 1 (p value 0.048), and CA associated to methylation cluster 3 (p value 0.000146). We identified enrichment of Wnt signaling genes in both AA and CA, however they were differentially activated in terms of canonical and non-canonical Wnt signaling pathway activation. This was further validated using the GenomeDx expression data. Our RRBS data also suggested distinct methylation patterns in AA compared to CA, and in part validated our TCGA findings. Survival analysis using the RRBS data suggested hypomethylated genes to be significantly associated with recurrence of prostate cancer in CA (p=6.07×10^−6^) as well as in AA (p=0.0077). Overall, the observed racial disparity in the molecular mechanism involved in the pathogenesis of prostate cancer suggests diverse heterogeneity that potentially could affect survival and should be considered during prognosis and treatment.

## Introduction

Prostate cancer (PCa) is one of the most prevalent malignancies affecting men (Fitzmaurice et al., 2013) [1] and the second-most leading cause of cancer-related death for men in the US [2,3]. According to Surveillance, Epidemiology, and End Results (SEER) there is a decline in prostate cancer incidence over the past few decades [4]. However, African American men (AA) present early onset of the disease with high-grade tumor, aggressive metastasis potential with increased risk of recurrence, and have worse survival outcomes when compared to their CA counterparts [5–8]. It is suggested that racial disparity in PCa is influenced by socioeconomic status, diet and lifestyle [9–11], but several lines of evidence indicate that biological factors like genetic and epigenetic factors are major contributors to this disparity [8, 12]. Such factors include ERG rearrangement, PTEN deletion, SPINK1 overexpression, SPOP mutation and 8q24 SNPs [13, 14]. Furthermore, PCa in AA is associated with higher testosterone levels and prominent immune related tumor biology [15–17], with tumor microenvironment dynamics associated with more aggressive phenotype in AA when compared to CA [18].

Aberrant DNA methylation is an important epigenetic modulation that has been implicated in PCa etiology and disease progression [19]. Methylation changes primarily occur in CpG islands sequestered in the promoter regions [20]. Whole genome methylation studies have identified a large number of hypo and hypermethylated regions in PCa patients [21, 22]. However, previous methylation-based racial disparity studies are limited and have either focused only on a specific gene or benign tissues [23, 24]. More recently, a methylation study using arrays reported differential methylation changes in ABCG5, MST1R, SNRPN and SHANK2 genes in AA and CA PCa patients [25]. They observed ABCG5 and SNRPN genes to be associated with the proliferation and invasion in cell lines of CA PCa patients [25].

In this study, we investigated the epigenetic variations in AA and CA PCa patients and how they are associated with survival and recurrence outcomes using the cancer genome atlas (TCGA) dataset [26] and Reduced Representation Bisulfite Sequencing (RRBS).

## Materials and Methods

### Prostate Tissue Samples

Prostate tumor tissues and adjacent benign prostate tissues were collected from patients at Icahn School of Medicine at Mount Sinai as per Institutional Review Board approved protocols (GCO#06-0996, 14-0318 and surgical consent). The fresh frozen prostatectomy specimens were procured after the tumor and benign regions were identified and marked by the pathologist. Three AA and three CA PCa patients were enrolled in this study, their clinical profile including tumor stage and Gleason score has been elaborated in **Table 1**. Genomic DNA was extracted from frozen tissues using Qiagen’s DNA/RNA extraction kit (Cat No. 80204) following the manufacturer’s instruction.

**Table 1.** Clinical profile of Prostate cancer patients.

| **Patient ID** | **Race** | **Gleason Score** | **PSA** | **Stage** |
| --- | --- | --- | --- | --- |
| CA1 | Caucasian | 4 + 5 | 15.5 | pT3b |
| CA2 | Caucasian | 4 + 3 | 6 | pT2c |
| CA3 | Caucasian | 5 + 4 | 1.5 | pT3a |
| AA1 | African American | 3 + 4 | 20.7 | pT3b |
| AA2 | African American | 3 + 4 | 4.8 | pT2c |
| AA3 | African American | 3 + 4 | 10 | pT3b |

### Reduced Representation Bisulphite Sequencing

Library preparation for RRBS was performed using NuGEN Ovation^®^ RRBS Methyl-Seq System (Cat N0: 0353,0553) for Illumina Sequencing. In brief, the preparation steps included: 1) DNA fragmentation by MspI enzyme digestion; 2) ligation of adapter; 3) end repair; 4) DNA oxidation; 5) Bisulphite conversion; 6) Desulfonation and purification; and 7) Amplification and purification. The libraries were sequenced using Illumina HiSeq and MiSeq platform (75bp paired end) at The Genomics Core Facility at the Icahn Institute and Department of Genetics and Genomic Sciences.

### Computational analysis

Bisulphite sequencing data was analyzed using Bismark v0.15.0 software. It is a set of tools for the Bisulfite-Seq data which performs alignments of bisulfite-treated reads to a reference genome as well as cytosine methylation calls at the same time. Bisulphite converted hg19 reference genome was used for the alignment and mapping of reads. Sequence reads that produce a unique alignment against the bisulfite genomes are then compared to the normal genomic sequence and the methylation state of all cytosine positions in the read is inferred. Read alignment details for RRBS data is summarized in **Table 2**. The methylation status of the uniquely mapped reads was extracted using the bismark_methylation_extractor script in Bismark. Only CpGs with at least five reads covering them were used for downstream analysis. Methylation of a specific region was calculated by averaging the methylation levels of all CpGs covered in that region. We calculated promoter methylation levels for each sample then compared the variation among CA vs AA and tumor vs benign region for each racial group. Promoters were defined as ± 2kb windows centered on RefSeq transcription start sites.

**Table 2.** Reduced representation bisulfite sequencing data information for prostate tissues (benign and tumor) of African American men and Caucasian men.

| **Race/**<br>**Group** | **Type of**<br>**tissue** | **Raw reads** | **Unique**<br>**mapped reads** | **Mapping**<br>**percentage** |
| --- | --- | --- | --- | --- |
| CA1 | Benign | 12282536 | 5416647 | 44.1% |
|  | Tumor | 12732930 | 5803964 | 45.6% |
| CA2 | Benign | 10462111 | 4675127 | 44.7% |
|  | Tumor | 10607889 | 4840047 | 45.6% |
| CA3 | Benign | 7160756 | 3072980 | 42.9% |
|  | Tumor | 11087471 | 5098151 | 46.0% |
| AA1 | Benign | 12788974 | 5762829 | 45.1% |
|  | Tumor | 13354714 | 5831607 | 43.7% |
| AA2 | Benign | 7520582 | 3363084 | 44.7% |
|  | Tumor | 12431991 | 5216656 | 42.0% |
| AA3 | Benign | 10402284 | 4513597 | 43.4% |
|  | Tumor | 9577045 | 3958823 | 41.3% |

### Survival analysis

SurvExpress analysis tool (http://bioinformatica.mty.itesm.mx:8080/Biomatec/SurvivaX.jsp) was used to determine the association between hypermethylated genes and survival. This tool uses both the gene list from our dataset and publically available human cancer database to predict biomarkers for risk assessment and survival. PRAD TCGA database [26] was used for the identification of biomarkers associated with survival. The Sobner [27] and Tyalor databases [28] were used for the analysis of survival and recurrence. The software generates risk groups by either by splitting the ordered Prognostic Index (risk score estimated by beta coefficients multiplied by gene expression values) for each samples at the median where samples are distributed equally in both groups or by using an optimized algorithm from the ordered prognostic index [29]. The survexpress software was used in default setting and it performs a log-rank test along all values of the arranged PI and then, the algorithm selects the split point where the p-value is minimum. This procedure is repeated until no changes are observed. This tool presents the output in the form of Kaplan-Meier curve for risk groups, concordance index which estimate the probability that subjects with a higher risk will experience the event after subjects with a lower risk and p-value of the log-rank testing equality of survival curves.

### TCGA data re-analysis

The Cancer Genome Atlas Network (2015) performed extensive genomic, transcriptomic, epigenetic and proteomics studies on a large cohort of prostate cancer patients (333 primary prostate tumor tissues) to identify the molecular and biological heterogeneity in prostate cancer patients [26]. In order to investigate the epigenetic variation and specific methylation pattern in AA and CA men, we retrieved the TCGA datasets with methylation cluster information and clinical details for each patient from the cBioPortal for Cancer Genomics (http://www.cbioportal.org/study.do?cancer_study_id=prad_tcga_pub). Of the 333 prostate cancer patients who were enrolled in TCGA study, 43 PCa patients were reported to be AA and 162 were CA. Further, to identify the epigenetic changes existing in each racial group we used gene lists that were reported to be dysregulated in each methylation cluster in TCGA study. The list of differentially regulated genes in each cluster of TCGA study was procured from firebrowse (http://firebrowse.org/?cohort=PRAD) [30]. We also used TCGA methylated data list to validate our RRBS analysis, which was retrieved from (http://firebrowse.org/?cohort=PRAD)portal [31].

### Functional analysis

The Ingenuity Pathway Analysis (IPA) software (Ingenuity Systems, Redwood City, CA) was used to identify the relevant bio-functions and the pathways associated with differentially expressed gene in each methylation cluster. The software generates pathways utilizing the list of genes from TCGA data and the list of genes stored in the ingenuity knowledge database, which is based on functional annotations and experimental observations. It also computes a p value for each pathway, to assess the likelihood of the association between the focus genes and a canonical pathway being not random. IPA also predicts the activation or inhibition of a canonical pathway based on z-score, which is automatically calculated based on differentially expressed genes in both the uploaded dataset and the information stored in IPA knowledge database. A positive z-score suggests the activation, whereas a negative z-score indicates inactivation of the pathway.

### GenomeDx analysis

Genomics Resource Information Database (GRID) patients’ samples were used for the validation of our observation of differential Wnt signaling in CA and AA cohorts from TCGA data. RNA was extracted from routine formalin-fixed, paraffin embedded (FFPE) Radical prostatectomy tumor tissues, which were amplified and hybridized to Human Exon 1.0 ST microarrays (Thermo-Fisher, Carlsbad, CA). All GRID data were normalized utilizing the single channel array normalization algorithm [32]. GRID samples were processed as described in [33].

## Results

### Identification of differential methylation clusters among AA and CA PCa patients in TCGA dataset

An integrative epigenetic and genetics analysis in TCGA study identified four distinct methylation groups which are named as methylation cluster (MC) 1, 2, 3 and 4 [26]. Prevalent PCa genetic alterations like ERG, ETV1, ETV4 fusions and SPOP, FOXA1, IDH1 mutations were reported to be differentially associated with each methylation clusters [26]. Interestingly, ERG fusion-positive PCa patients were differentially distributed in MC1 and MC3; one-third of them exhibited hypermethylation and belonged to methylation cluster (MC1) whereas, two-third of ERG positive PCa patients had hypomethylated DNA loci (MC3) **Figure 1A**. MC2 had patients with SPOP, FOXA1 and IDH1 mutations. They also included ETV1 and ETV4 fusion and had highly methylated DNA. MC4 was positive for ERG, ETV1 and ETV4 fusion as well as SPOP and FOXA1 mutations.

**Figure 1.**
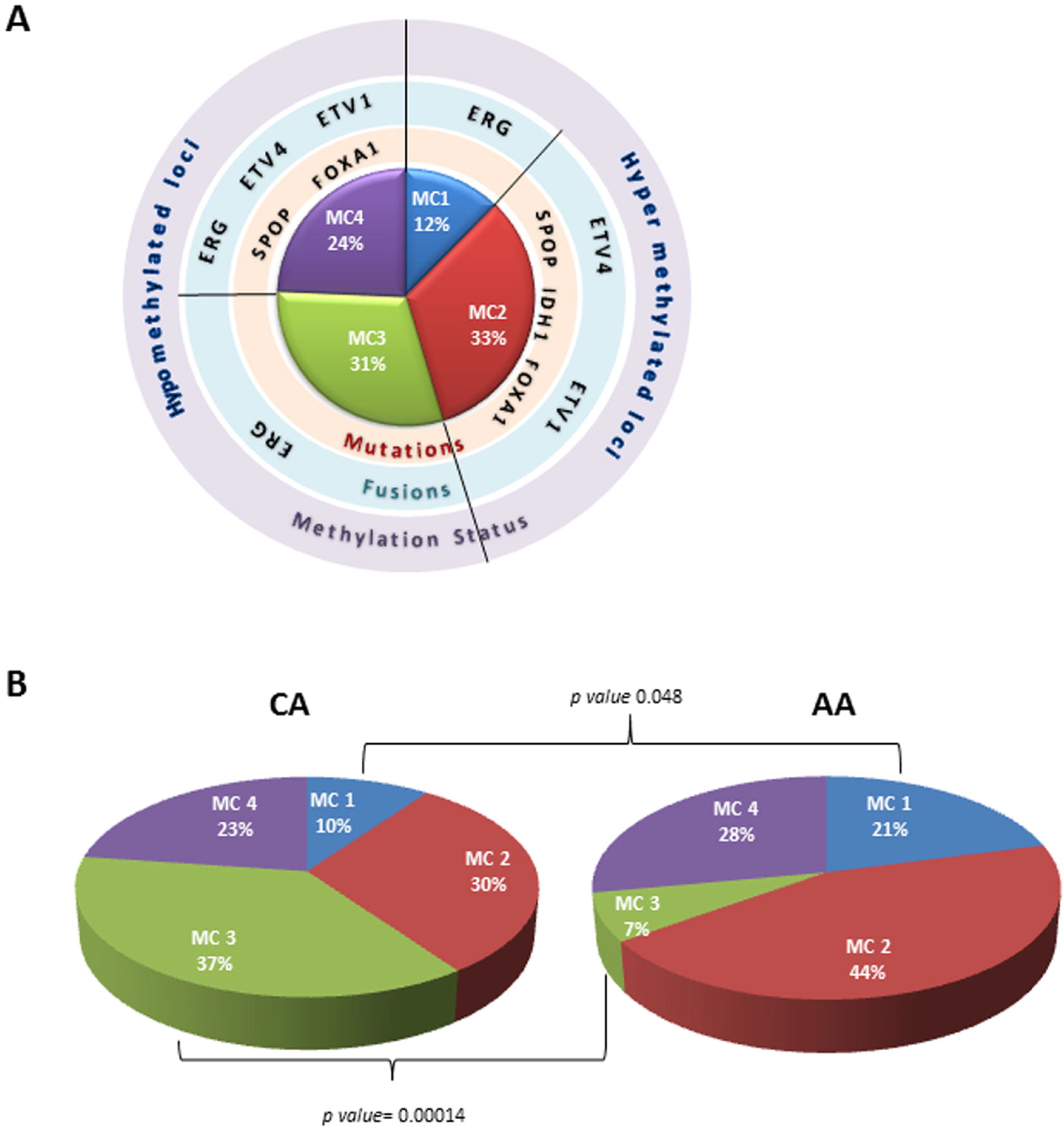
A. Pictorial representation for the distribution of fusion and mutation associated tumors in different methylation clusters. B. The distribution of African-American and Caucasian patients in different class of methylation clusters. African-American patients are more likely to belong to MC1 whereas Caucasian preferentially belongs to MC3.

In this study, we re-analyzed TCGA data for identification of epigenetic variation among AA and CA racial groups. PCa patients where then divided into two groups based of their race (43 AA and 162 CA) and we observed that AA men were more likely to present the MC1 pattern (p value 0.048), while CA men mostly presented with MC3 (p value 0.000146), **Figure 1B**. The MC1 cluster displays a distinct spectrum of hypermethylated loci whereas the MC3 cluster shows hypomethylated loci.

### Pathways associated with different methylation clusters

To study the predominant signaling pathways associated in each methylation cluster we retrieved the mRNA expression data from TCGA and generated a list of differentially expressed gene in each methylation cluster. We used the ingenuity pathway analysis (IPA) to identify pathways affected in each methylation cluster. We observed Wnt signaling pathway to be differentially regulated in all four clusters suggesting it to play an important role in PCa pathogenesis **(Supplementary Table 1)**. Since AA and CA PCa patients distributed distinctly in MC1 and MC3 only we focused our analysis on these two clusters only. The Analysis of MC1 showed that non-canonical Wnt signaling (Wnt/Ca^+2^ signaling) was downregulated in CA whereas it was activated in AA, **Figure 2A**. The analysis of MC3 showed exclusive activation of non-canonical Wnt/Ca^+2^ in CA. Furthermore, inflammation/immune cell related pathways and PI3K signaling was observed to be associated with upregulated genes in MC1 (whereas down regulated genes associated with cell cycle signaling). In the case of MC3, amino acid metabolism related pathways were associated with upregulated genes, and downregulated genes were associated with cytokine signaling and PI3K-pathway alterations **Table 3**.

**Figure 2.**
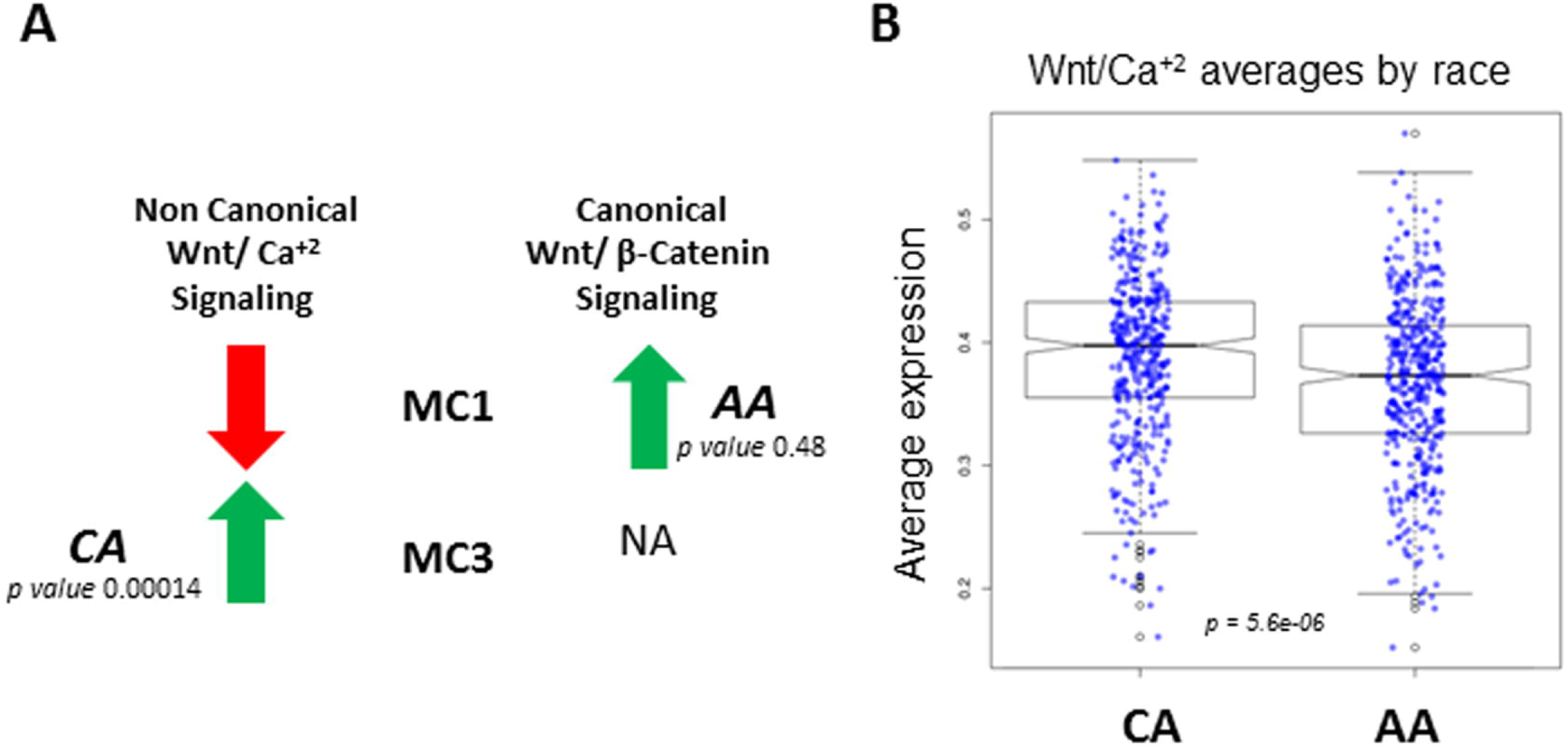
A. Pictorial representation of differential regulation of canonical (Wnt/β-catenin) and non-canonical (Wnt/Ca^+2^) Wnt signaling pathways among methylation cluster 1 and 3. B. Wnt (Wnt/Ca^+2^) signaling genes were significantly overexpressed in CA cohort as compared to AA cohort from GRID data from GenomeDx.

**Table 3.** Pathways associated with differentially expressed genes in methylation cluster 1 and 3.

|  | **Methylation cluster 1**<br>**(African-American)** |  | **Methylation cluster 3**<br>**(Caucasian)** |  |
| --- | --- | --- | --- | --- |
|  | **Pathways** | **Genes** | **Pathways** | **Genes** |
| **Upregulated** | Chemokine signaling and Immune cell or inflammation related signaling | MCP-3, Eotaxin, PKC, MLCP, CCR3, PI3K, cPLA2, Gβ, AIM2, CASP1, CASP5, VAV2, PXN, VCAM1, PRKCQ, NOX3, MMP16, CLDN18, PRKCZ, CTNNA2, IRS2, DLC1, MMP17, ITK, PRKCB | Amino Acid Metabolism | BCAT1, BCAT2, HMGCLL1, EHHADH, ALDH2, ALDH1A2, ALDH1L1, ALDH3A1 |
|  | PI3K Signaling | VAV2, PLCZ1, PTPRC, IRS2, PLCL2, PRKCZ, NFATC1, PPP3CA, PRKCB |  |  |
| **Downregulated** | Cell Cycle related pathways | CDKN2A, HDAC9, CCND2, CCNA1, CCND3, PPP2R2A, PPP2R3A, HDAC7, CUL1, CDK6, HDAC10, PPP2R5C, CDKN1B, NRG1, SMAD3 | Cytokine and Chemokine signaling | VCAM1, IL-6, AKT, PI3K, JAK1, JAK3, PLCγ1, mTOR, NFAT, MHC-II, MCP-3, SDF-1, CAMK, PI3Kγ, MLCP, CCR3, |
|  |  |  | PI3K Signaling | NFAT, TAPP, PLCγ1, CAMK, IP3R, Calcineurin, TLR4, PI3K, AKT |

The finding of differential regulation of Wnt signaling in MC1 (representation of AA cohort) and MC3 (representation of CA cohort) from TCGA data re-analysis were further validated in a separate and larger cohort of PCa patients among these two racial groups. We used the GRID data from GenomeDx, which uses RNA expression analysis from prostate tumors to provide a Decipher^R^ genomic score that predicts disease aggressiveness. In this data base, genes belonging to the non-canonical Wnt/Ca^+2^ signaling pathway were significantly upregulated in CA as compared to AA (p value 5.6×10^−06^), **Figure 2B**.

### Epigenetic variation in AA and CA Prostate Cancer patients

To further validate our TCGA analysis and identify the epigenetic differences in AA and CA prostate tumors, we performed RRBS analysis on 6 PCa patients, three in each group, **Table 1**. Benign tissue adjacent to tumor was procured from each patient to serve as control. We compared the methylation changes in CA and AA tumor tissues and observed very distinct methylation pattern in both the racial groups. Further, analysis revealed hypermethylation of 33 genes in AA, 19 hypermethylated genes in CA (**Supplementary table 2)**. This suggests hypermethylation to be a predominant event in AA, which is in concordance with our observation with TCGA data where MC1 cluster had more hypermethylated genes compared to MC3 **Figure 3A**. Moreover, we observed significant methylation changes in benign vs tumor tissue in both CA and AA **Figure 3B and C**, respectively.

**Figure 3.**
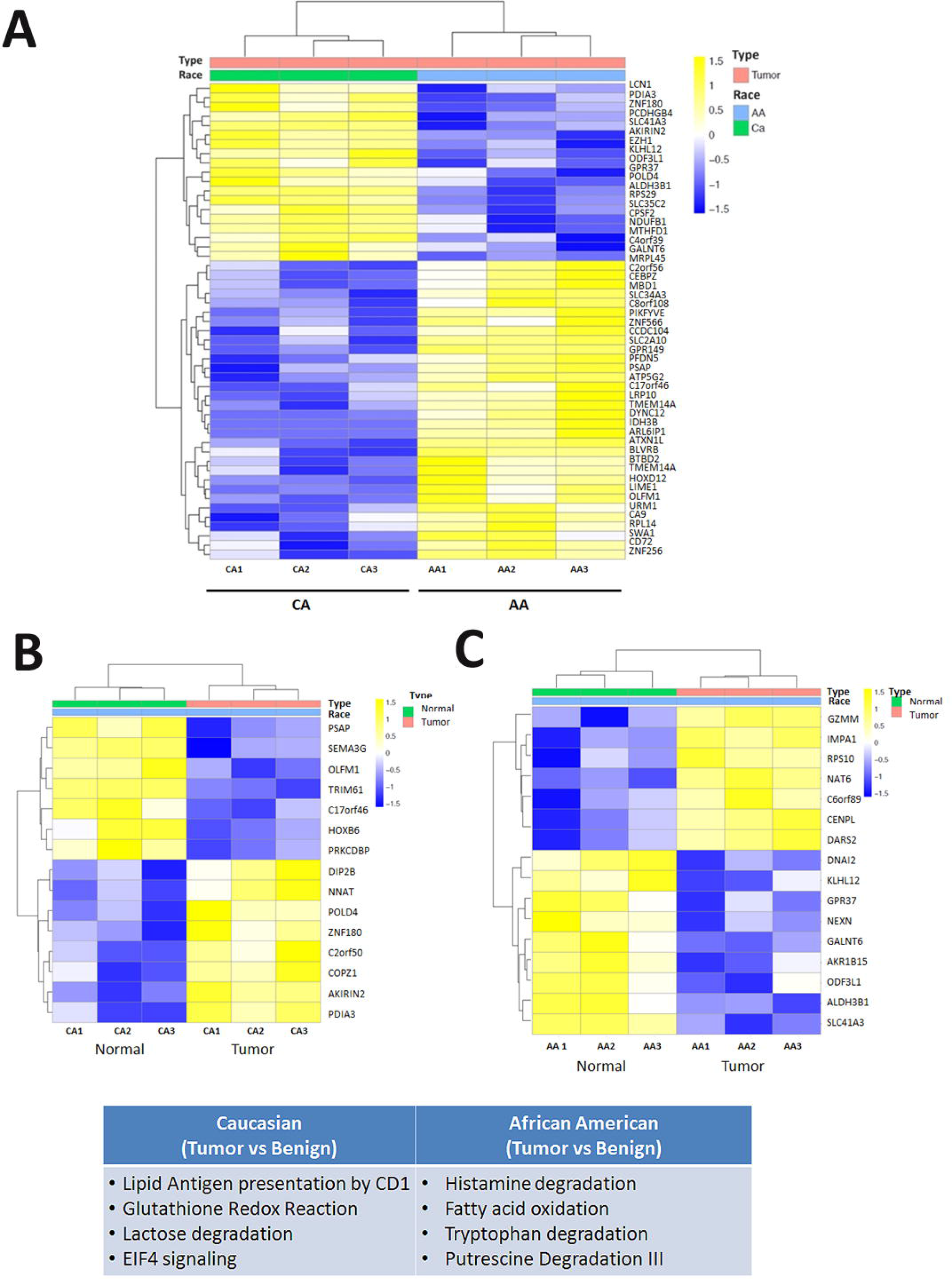
Epigenetic changes in Caucasian and African-American Prostate cancer patients. A. Heat map representing the methylation variation in CA and AA. B. Heat map represents the methylation variation in benign vs tumor tissues in CA. C. Heat map represents the methylation variation in benign vs tumor tissues in AA Yellow color correspond to hypermethylated genes in both the groups whereas blue color represents the hypomethylated genes.

### Correlation of RRBS analysis with TCGA data

To validate our finding we matched the methylated gene list from RRBS with the TCGA methylation data. We used the negative correlation between DNA methylation and mRNA expression data from TCGA for this comparison [31]. Methylation mean with cut-off < 0.2 was used for hypomethylated genes and cut-off > 0.3 was used for hypermethylated genes [26]. Even though different platforms were used for the methylation studies, in both the cases, we observed similar methylation patterns; RPS29, ZNF180, POLD4, AKIRIN2, SLC41A3, PDIA3, CPSF2, and SLC35C2 were found to be hypomethylated in AA men in RRBS analysis as well as in TCGA analysis, **Table 4**. Hypermethylated genes from both the analysis include IDH3B, LRP10, BLVRB, ZNF256, LIME1, CD72, URM1, PSAP, PIKFYVE, ARL6IP1 and BTBD2, **Table 5**.

**Table 4.** Comparison of hypomethylated genes in African American (hypermethylated in Caucasian) in RRBS analysis and hypomethylated in TCGA dataset (Methylation mean cut off < 0.2).

|  | **RRBS** |  | **TCGA data** |  |
| --- | --- | --- | --- | --- |
| **Gene** | **P value** | **Methylation difference** | **P value** | **Methylation Mean** |
| RPS29 | 0.002 | -0.23 | 1.07E-09 | 0.04 |
| ZNF180 | 0.03 | -0.15 | 1.04E-02 | 0.03 |
| POLD4 | 0.04 | -0.14 | 3.58E-06 | 0.14 |
| AKIRIN2 | 0.01 | -0.13 | 4.67E-15 | 0.03 |
| SLC41A3 | 0.02 | -0.13 | 4.69E-03 | 0.06 |
| PDIA3 | 0.01 | -0.12 | 7.33E-02 | 0.12 |
| CPSF2 | 0.04 | -0.11 | 1.62E-07 | 0.04 |
| SLC35C2 | 0.01 | -0.10 | 5.60E-08 | 0.05 |

**Table 5.** Comparison of hypermethylated genes in African American (hypomethylated in Caucasian) in RRBS analysis and hypermethylated in TCGA dataset (Methylation mean cut-off > 0.3)

|  | **RRBS** |  | **TCGA data** |  |
| --- | --- | --- | --- | --- |
| **Gene** | **P value** | **Methylation difference** | **P value** | **Methylation Mean** |
| IDH3B | 0.035 | 0.262 | 9.95E-09 | 10.38 |
| LRP10 | 0.017 | 0.105 | 1.66E-04 | 12.16 |
| BLVRB | 0.039 | 0.166 | 7.89E-03 | 10.45 |
| ZNF256 | 0.021 | 0.135 | 1.02E-05 | 6.71 |
| LIME1 | 0.047 | 0.205 | 3.03E-07 | 6.34 |
| CD72 | 0.038 | 0.106 | 4.15E-15 | 4.47 |
| URM1 | 0.017 | 0.266 | 1.49E-03 | 10.25 |
| PSAP | 0.008 | 0.467 | 5.37E-09 | 14.88 |
| PIKFYVE | 0.046 | 0.106 | 1.12E-06 | 9.96 |
| ARL6IP1 | 0.003 | 0.202 | 3.14E-15 | 12.12 |
| BTBD2 | 0.032 | 0.130 | 6.64E-07 | 11.49 |

### Correlation of hypomethylated genes with survival in AA and CA Prostate cancer patients

To identify whether the genes we identified had any prognostic significance, we performed survival analysis on the hypomethylated genes obtained from RRBS analysis in both the racial groups. PRAD-TCGA (TCGA) dataset (497 samples) [26] and Sboner datasets (281 samples) [27] were used for the prediction of survival outcome whereas Taylor datasets (140 Sample) [28] was used to access the recurrence of prostate cancer in different racial group. Both Sboner data and TCGA data were able to separate the risk group based on differential gene expression listed in **Table 6**. Nonetheless, p value for the risk group separation was better for Sboner dataset as compared to TCGA data for both the racial groups (**Figure 4A, B**, **Table 6**). In addition, Taylor datasets suggested significant association of hypomethylated genes with biochemical recurrence of prostate cancer in CA (p value 6×10^−6^) as well as in AA (p value 0.0077) (**Figure 4C**, **Table 6**).

**Figure 4.**
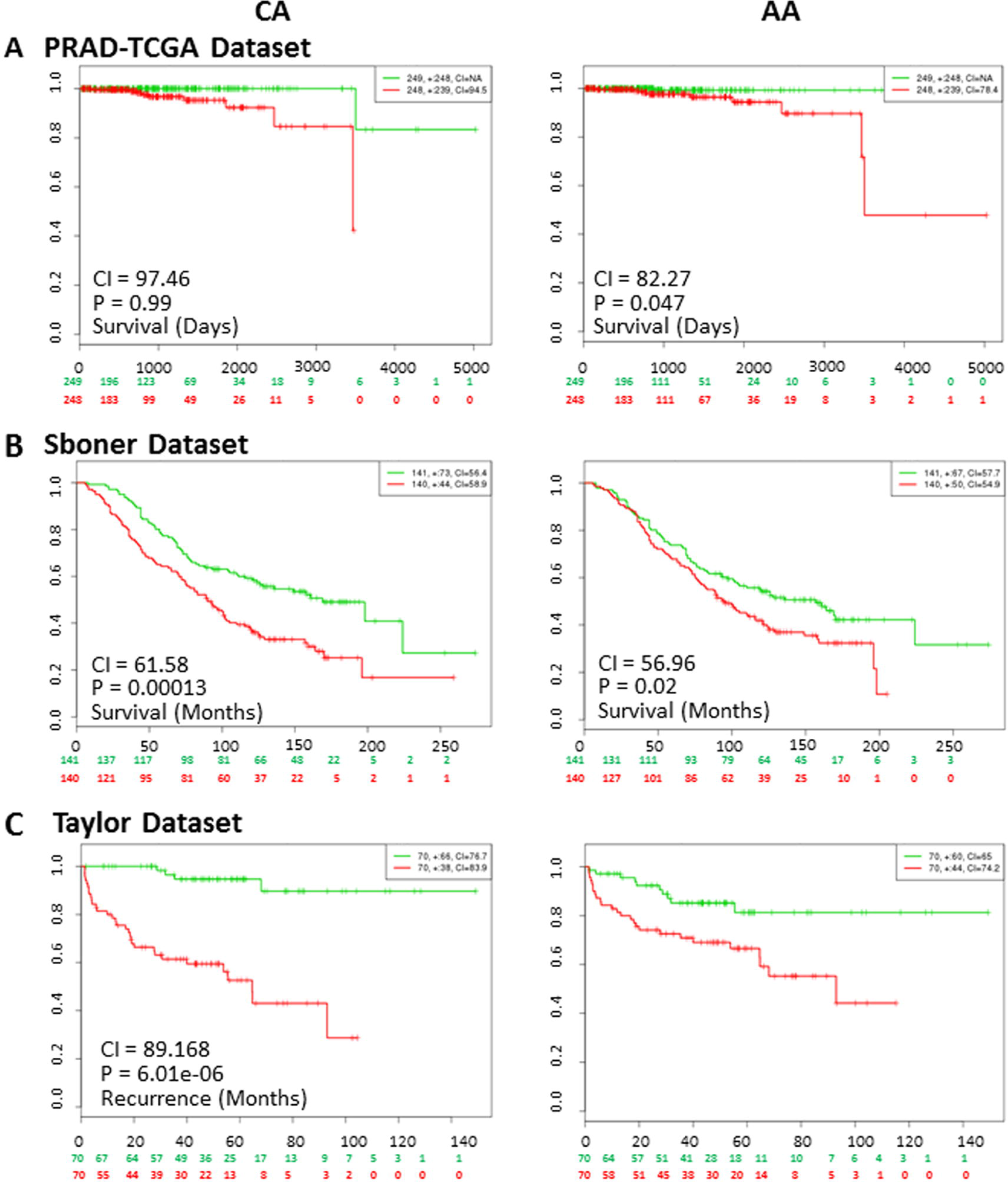
Comparison of Kaplan--Meier curve in Caucasian and African American Prostate cancer patients using three prostate cancer databases using survexpress software. A. PRAD-TCGA prostate adenocarcinoma database shows survival days. B. Sboner prostate database survival months. C. Taylor prostate database shows recurrence months. Horizontal axis represents time to event (days in A, months in B and C). Red and Green curves denote High-and Low-risk groups respectively. The red and green numbers below horizontal axis represent the number of individuals not presenting the event of the corresponding risk group along time. The number of individuals, the number of censored, and the CI of each risk group are shown in the top-right insets. CI represent Concordance index, P represent P value.

**Table 6.** Evaluation of survival analysis and prognostic markers in three different prostate cancer databases.

| Datasets | Caucasian |  |  | African American |  |  |
| --- | --- | --- | --- | --- | --- | --- |
|  | Risk group p Value | DEG between risk groups | Genes | Risk group p Value | DEG between risk groups | Genes |
| **PRAD-TCGA database [26]** | 0.998 | 14 | HOXD12, C6orf108, C17orf46, SIVA1, OLFM1, TMEM14A, C2orf56, BLVRB, CA9, ZNF566, ARL6IP1, CCDC104, MBD1, PSAP | 0.0474 | 12 | KLHL12, EZH1, RPS29, ALDH3B1, LCN1, MTHFD1, GALNT6, AKIRIN2, ODF3L1, NDUFB1, SLC35C2, GPR37 |
| **Sboner database [27]** | 0.00013 | 10 | ATP5G2, C6orf108, CD72, SIVA1, OLFM1, CEBPZ, BLVRB, CA9, IDH3B, PSAP | P = 0.02 | 7 | EZH1, ALDH3B1, LCN1, MTHFD1, POLD4, PDIA3, GPR37 |
| **Taylor database [28]** | 6.01e-06 | 10 | ATP5G2, SLC34A3, PIKFYVE, ATXN1L, DYNC1I2, CEBPZ, CA9, ARL6IP1, URM1, MBD1 | P = 0.0077 | 8 | EZH1, LCN1, GALNT6, POLD4, C4orf39, SLC41A3, ODF3L1, CPSF2 |

## Discussion

Comprehensive “omics” studies of Prostate cancer in the cancer genome atlas (TCGA) study indicated that diverse heterogeneity exists in PCa patients [26]. Besides, this fact there exist disproportionate incidence, progression and mortality of PCa patients in relation to race. In this study we identified that AA and CA have distinct methylation patterns. Further, pathway analysis revealed that the specific methylation events in each group are associated with different biological functions. Each racial group is associated with differential Wnt signaling and other different biological pathways, which have not been explored in details previously with respect to racial disparity. We also observed discrete methylation changes of genes in AA and CA patients which may contribute to differences in prostate cancer outcomes between these two populations. Epigenetic changes are an inherent genomic property that appears early during development [34] or sometime acquired during lifetime due to exposure of other environmental factors [35, 36]. Thus it is a critical factor in the pathogenesis of PCa and could contribute towards aggressive clinical outcomes.

Several studies reported that hypermethylation of CpG is the predominant event occurring in case of prostate cancer [26, 37]. There are large variations in methylation of specific genes among AA and CA ethnic groups [22–24], reports suggest the existence of race specific methylation differences at the birth [36, 38]. Using the TCGA data and RRBS analysis, we observed that AA have predominant methylation events as compared to CA, and both the groups have very distinct spectrum of methylated loci. This result of race specific global methylation difference is interesting and requires further investigation to elucidate the mechanism driving this disparity.

Further, pathway analysis also suggests disparity in terms of biological functions and signaling mechanism which is a crucial event. For instance, we found canonical and non-canonical Wnt signaling pathways are differentially regulated in AA (MC1) and CA (MC3). Wnt/β-catenin signaling is upregulated and Wnt/Ca^+2^ downregulated in AA whereas; the later is downregulated in CA. Although, multiple studies have identified association of Wnt signaling with aggressive, late stage disease, and metastasis in prostate cancer [39–41], to our knowledge, the distinction of Wnt/β-catenin and Wnt/Ca^+2^ signaling pathway association with ethnicity in prostate cancer has not been reported previously. Of the two Wnt signaling pathways, Wnt/β-catenin is very well studied pathway and plays important role in progression of prostate cancer towards metastasis, and development of disease [42, 43]). More recently, non-canonical Wnt (Wnt/Ca^+2^) signaling has also been implicated in the pathogenesis of prostate cancer [41, 44]. The involvement of Wnt/β-catenin pathway in progression could be attributed to its function in proliferation and survival of cancer stem cells and was suggested to act as molecular markers for predicting resistance to several androgen dependent therapies [45]. Lately, genes involved in canonical and non-canonical Wnt signaling pathways are emerging as promising targets for therapeutic intervention in prostate cancer [46, 47]. It would be interesting to explore its therapeutic role in prostate cancer patients of different ethnicities.

We also observed that inflammation related pathway, PI3K signaling, cell cycle signaling, and amino acid metabolism pathways vary differentially in these racial groups. Previous studies have suggested association of interleukins signaling and inflammation to be more prevalent in AA relative to CA men [16, 48]. Also, PI3K signaling is crucial in PCa as Phosphatase and tensin homologue (PTEN) loss, resulting in constitutive activation of the PI3K pathway, is common in advanced prostate cancer. Using reciprocal-pairing of miRNA-mRNA, Wang et al., 2015 predicted the overt activation of PI3K/AKT pathway in African American men compared to CA men [49]. Our observations are similar to these findings. Furthermore, we found cell cycle signaling was downregulated in AAs whereas a recent study shows enhanced cell cycle related transcription in specific subpopulation of CA derived prostate cancer cell-line [50]. Finally, similar to our observation of amino-acid metabolism to be associated with CA, Putluri et al., 2011 reported higher amino acid metabolism in CA derived prostate cancer cell lines treated with androgens [51].

From the present study we also identified distinct set of methylated genes in AA and CA men using RRBS. We observed genes hypermethylated in AA were hypomethylated in CA and vice-versa. Some genes were also found to follow the same methylation pattern as observed in TCGA study which validates our finding. These include the PDIA3 gene which was hypomethylated in AA (hypermethylated in Ca) as well as in TCGA data. Evidence suggests its association with malignant stage of prostate cancer which could be used as potential biomarkers [52]. Furthermore, hypermethylated genes in AA include isocitrate dehydrogenase beta-subunit (IDH3B) and Phosphoinositide Kinase, FYVE-Type Zinc Finger Containing (PIKFYVE), which a study suggests could serve as a urine-based biomarker for prostate cancer [53]. The prostate has been suggested to synthesize high amount of citrate and mutations in isocitrate dehydrogenases have been associated with PCa [54]. The other gene PIKFYVE induces exosome secretion and is known to influence cancer related pathways like metastasis and drug resistance development. [55]. However, the importance of these genes with respect to racial disparity in PCa needs to be explored further.

Using in-silico analysis we observed that the Sboner dataset [27] and Tylor datasets [28] predicted survival and recurrence outcome in both the ethnic group with the RRBS data. However, lack of ethnic information in the available dataset limits the predicting impact of these genes in specific ethnic populations. Thus, study involving large cohort of AA and CA Prostate cancer patient is needed for the better prediction of bio-markers for risk assessment, aggressive cancer prediction recurrence and survival outcome which will eventually facilitate clinical translation of genome-based prognostic biomarkers.

This is an era of personalized treatment and genomic, epigenomic and proteomic approaches has contributed a lot in the stratification of various complexities with respect to active surveillance, surgery, therapeutics in prostate cancer. The present study indicates that prostate cancer patient of different ethnicity have distinct epigenetic alterations and biological functions, and thus should be managed differently. Further, study with larger cohort of AA and CA PCa is needed with detailed clinical information to predict better biomarkers for the outcomes and for the targeted therapeutic interventions.

## Acknowledgements

We are thankful to the Department of Hematology and Oncology, Icahn School of Medicine at Mount Sinai, New York for use of their equipment and resources. Both SSY and KKY thank the Prostate Cancer Foundation for Young Investigator Awards.

